# Flower visitation of alien plants is non-linearly related to phylogenetic and floral similarity to native plants

**DOI:** 10.1101/2022.02.14.480304

**Authors:** Mialy Razanajatovo, Felana Rakoto Joseph, Princy Rajaonarivelo Andrianina, Mark van Kleunen

## Abstract

1. Biological invasions are key to understanding major ecological processes that determine the formation of novel interactions. Flower visitation to alien species may be facilitated by co-flowering natives if they share similar floral traits with the latter. However, if competition for pollinators is important, flower visitation to alien species may be higher if they have traits different from those of native species. We tested whether flower visitation to alien plants depended on phylogenetic relatedness and floral similarity to native plants.
2. In a field experiment, we simulated invaded communities by adding potted alien plants into co-flowering native communities. We recorded flower visitation to pairs of 34 alien and 20 native species totalling 151 species combinations for 3,068 hours.
3. Flower visitation to alien species was highest when they had intermediate floral trait distances to native species, and either low or high phylogenetic distances. The alien plants received more similar flower-visitor groups to natives when they had low phylogenetic and either low or high floral trait distances to native plants.
4. The non-linear patterns between flower visitation and similarity of the alien and native species suggest that an interplay of facilitation and competition simultaneously drives the formation of novel plant-pollinator interactions. The shapes of the relationships of phylogenetic and floral trait distances with flower visitation to alien plants were contrasting, possibly due to different strengths of phylogenetic signal among traits.
5. We provide experimental evidence for the effects of relatedness and functional similarity to native plants on flower visitation of alien plants. We show that such effects might be non-linear, and that effects of trait dissimilarity and phylogenetic distance on pollinator-mediated interactions can reflect different mechanisms.

## Introduction

Biological invasions are a major characteristic of human-induced global environmental change. Invaded ecosystems and communities in many parts of the world have been affected severely (Fei et al., 2014; Vilà & Hulme, 2017). For this reason, much of the efforts in ecological research has aimed at understanding what determines invasiveness. Furthermore, biological invasions represent natural experiments that offer real-time opportunities to study the assembly of communities (Shea & Chesson, 2002; Tilman, 2004). Specifically, as many alien organisms have integrated into native resident communities, biological invasions are key to understanding the major ecological processes that determine the formation of novel interactions.

As related species are likely to be more similar, they should show strong niche overlap and compete for shared resources. Based on this premise, Darwin (1859) posed that relatedness between the alien and the native species could impede the success of alien species (Darwin’s Naturalization hypothesis). At the same time, if there are related native species, this indicates that the environment will most likely be suitable for the alien species too (Preadaptation hypothesis). These two hypotheses, with opposing predictions on how relatedness between alien and native species should affect invasion success, are now known as Darwin’s Naturalization Conundrum (Diez et al., 2008). Indeed, the results of previous studies testing these hypotheses are largely inconsistent, suggesting that the two mechanisms act at different scales and stages of invasion (Cadotte et al., 2018; Thuiller et al., 2010). Moreover, most of the previous studies were correlative, based on floristic lists and field observations (Cadotte et al., 2018; Gallien & Carboni, 2017; Sheppard et al., 2018), whereas manipulative experiments, in which species are introduced into communities to provide causal insights (e.g. Malecore et al., 2019), are scarce. Furthermore, most previous studies considered direct interactions between alien and native plant species (e.g. competition for space or nutrients), but very few studies have tested indirect interactions such as those mediated by pollinators (but see Bezeng et al., 2015; Burns et al., 2011). However, as about half of all flowering plant species relies on pollinators for at least 80% of their seed production (Rodger et al., 2021), the ability to attract resident pollinators can play a major role in the integration of alien plants in novel communities.

To reproduce in the non-native range, alien plants, which are often decoupled from their historical pollinators can use resident pollinators (Razanajatovo et al., 2015; Razanajatovo & van Kleunen, 2016; Traveset & Richardson, 2014), and thereby form novel plant-pollinator interactions. There have been two apparently contradicting major concepts on the roles of plant-pollinator interactions in the assembly of invaded communities (Sargent & Ackerly, 2008). First, in the case of pollinator facilitation, alien plant species that can use the same pollinators as the native plant species (because they have more similar floral traits) should more readily attract pollinators, with similar abundance and composition to native plant species. Second, in the case of pollinator-mediated competition, alien plant species that can use pollinators different from those of native plant species should more readily attract pollinators in the local community. In both cases, the formation of novel plant-pollinator interactions between alien plants and resident pollinators should depend on the plant traits that influence flower visitation.

To advance our understanding of the ecological processes that govern the formation of novel plant-pollinator interactions, a trait-based approach (Kraft & Ackerly, 2014; McGill et al., 2006) has been suggested (Aslan et al., 2015). Floral traits, such as flower symmetry, color and size, can act as signals for flower visitors to locate floral rewards, and have therefore been considered as important traits that mediate plant-pollinator interactions (Fornoff et al., 2017; Junker et al., 2015; Ortiz et al., 2021; Reverté et al., 2016). Furthermore, nectar production has been shown to influence the indirect interactions between co-flowering plants with shared pollinators (Carvalheiro et al., 2014). Nevertheless, the floral traits involved in pollinator attraction are still generally missing from studies in plant-community ecology (E-Vojtkó et al., 2020; Sargent & Ackerly, 2008). A meta-analysis seeking to understand the impacts of alien species on the pollination and reproductive success of native species considered floral traits, and showed that similarities in flower symmetry and color in alien and native plants increased competition for pollinators (Morales & Traveset, 2009). Therefore, similarity in these floral traits may play critical roles in pollinator-mediated alien-native plant interactions.

The patterns of novel plant-pollinator interactions may differ with regard to phylogenetic and floral trait distances. While phylogenetic relatedness is frequently assumed to be a proxy for trait similarity, some floral traits may not be evolutionarily conserved (Sargent & Ackerly, 2008). For example, closely related species frequently have different flower colors (Eaton et al., 2012; Shrestha et al., 2014). Alien plant species that are phylogenetically closely related to native species may have either similar or dissimilar traits to native species. Also, alien species that are phylogenetically distantly related to native species may have either similar (convergent evolution) or dissimilar traits to native species. It is therefore important to consider both phylogenetic distance and trait dissimilarity (Cadotte et al., 2013; Lemoine et al., 2015), and to disentangle the effects of both (Marx et al., 2016). Thus, the patterns of novel plant-pollinator interactions might be more complex than those predicted by Darwin’s Naturalization Conundrum (Diez et al., 2008). Furthermore, different mechanisms may act simultaneously to drive the success of alien species, as has been shown for direct plant-plant interactions (Malecore et al., 2019; Sheppard et al., 2018). When both pollinator facilitation and competition for pollinators play a role, flower visitation to alien species should be highest or lowest at intermediate phylogenetic and floral trait distances to native species. It is therefore important to consider nonlinear relationships between flower visitation to alien species and phylogenetic and floral trait distances to native species.

In a field experiment in which we simulated invaded communities by adding potted alien plants into co-flowering native communities, we tested whether flower visitation to alien plants depended on phylogenetic relatedness and functional similarity to native plants. For pollinator facilitation and competition for flower visitors, respectively, we predicted a negative and a positive relationship between flower visitation to alien species and phylogenetic or floral trait distance to natives. If both mechanisms were operating, we predicted non-linear relationships with either high or low flower visitation at intermediate phylogenetic or floral trait distances (i.e. hump- or U-shaped relationships). Most previous studies on pollinator-mediated alien-native plant interactions investigated the impacts of co-flowering alien plants on the pollination and reproductive success of native species. Our study, in contrast, assessed the outcomes of pollinator facilitation by and competition for flower visitors with native species on alien species to understand how alien plants attract pollinators in the invaded range. More specifically, we asked (1) whether flower visitation to alien plants was related to the phylogenetic and floral trait distances between the alien and native species, and (2) whether the similarity in flower visitor composition was related to the phylogenetic and floral trait distances between the alien and native species.

## Materials and Methods

### Study species and sites

To simulate invaded communities in a field experiment in central Europe, we selected 34 herbaceous insect-pollinated neophytes (i.e. alien species introduced after the discovery of the Americas in 1492), covering a broad variation in floral traits such as size, symmetry and colors, occurring in semi-natural grasslands and anthropogenic or ruderal habitats, and usually flowering between April and September. The alien species belonged to 14 plant families, 68% were short-lived (annual or biennial), and 24% were self-incompatible (Supporting Information Table S1). The species neophyte status was based on information in the Floraweb (http://floraweb.de/index.html) and the Biolflor (Kühn et al., 2004) databases. We precultivated the alien plant species from seeds or seedlings (Table S1) in the research garden of the University of Konstanz in Germany (http://www.uni-konstanz.de/botanischergarten/) until they flowered. From May to September 2018, we carried out the field experiment in managed meadows near the city of Konstanz (Table S2).

### Experimental set-up and flower-visitation recording

Two to three days before adding alien plants to flowering resident native communities, we prospected grasslands around the city of Konstanz, and identified sites that were visibly dominated by one flowering native species, which served as the host native species. We then recorded the density of flowers as the number of flower units per 1m^2^ for all flowering species recorded in one 25 m^2^ plot at each site. To standardize the number of flower units across different species, we considered a flower unit to correspond to a receptacle area of 1 cm^2^ (Carvalheiro et al., 2014). We selected a total of 25 sites where the density of flowers for the host native species and for all flowering species ranged from 24 to 4,900 and from 10 to 1,763, respectively (Table S2). To simulate invaded communities, we placed for up to ten alien species (range=4-10, median=5) up to five (range= 2-5, median=4) potted flowering individuals into a site. The exact numbers depended on the availability of flowering alien plants. We paired each added alien plant individual with a host native plant individual. To let the alien plants adjust to the flower-visitor communities, and the insects to the newly added plants, we left the alien plants for two to three days at each site before we recorded flower visitation. We used a total of 20 host native species, i.e. the dominant flowering native species at a site (Table S2), belonging to ten plant families. In total, we had 151 combinations of added alien and host native species, spanning phylogenetic distances between the alien and the native species from 10.64 to 295.60 (median=236.40) million years (for the calculation of phylogenetic distance, see below).

To record flower visitation, we placed a BRINNO TLC200 time-lapse camera (https://www.brinno.com/time-lapse-camera/TLC200) at a vertical distance of 25-30 cm above the flowers of the paired flowering alien and native plants (Fig. S1). We set the time-lapse interval at 2 seconds and recorded from 10:30 to 16:30 on a sunny day (except for two set-ups in which we recorded from 9:30 to 14:30 due to logistic constraints). Sampling was done on one day for each setup. By using 40 cameras, we could observe many pairs of plants simultaneously. The alien plants were removed from the experimental plots at the end of the recording day. We collected a total of 3,068 hours of observations, which corresponded to 3 terabytes of video files. We analyzed each video file manually using the Blender software (https://www.blender.org/). The video analysis consisted of counting flower visits to the alien and the native plants in each species pair. We considered a flower visit when the flower visitor made contact with reproductive organs (anther and stigma). We attributed each flower visitor to one of the following flower visitor groups: Hymenoptera: honeybees (Apis spp.), bumblebees (Bombus spp.), other bees and wasps; Coleoptera (beetles); Diptera: hoverflies/syrphids and other flies; Lepidoptera: butterflies and moths; Mecoptera; Neuroptera; other unknown groups (impossible to identify). We recorded flower visitation for each one-hour interval within the whole recording time. We also counted the number of observed alien and native flower units within each frame.

### Measurements of floral traits

Using five plants per added alien and host native species at each site, we measured floral traits that most likely influence flower-visitor attraction. We recorded flower symmetry (radial or bilateral), and measured flower size as the diameter or the largest width of a flower. To measure flower color, we measured floral reflectance spectra using an AvaSpec-2048 fibre optic spectrometer and an Ava Light-XE xenon light source (Avantes, Eerbeek, The Netherlands) relative to a standard white reference tile (WS-2) at an angle of 90°. We measured the reflectance spectra of five corolla samples (each from different plants) per added alien and host native species. We classified the spectra into four binomial categories: blue (wavelength 401-470 nm), green (471-540 nm), yellow (541-610 nm) and red (611-680 nm), based on the presence/absence of local maxima at the respective wavelength interval (Fornoff et al., 2017). As a measure of the presence of floral rewards, we added data on nectar production (yes/no) using database and literature sources (Table S3).

### Calculation of phylogenetic and floral trait distances

To test whether flower visitation to the alien species was influenced by relatedness and floral similarity between the alien and the native species, we calculated phylogenetic and floral trait distances. We constructed a phylogenetic tree for the alien and native species by pruning the dated DaPhnE supertree (Durka & Michalski, 2012). Because *Hypochaeris radiata* was not included in the DaPhnE tree, we added a tip at the root of the Hypochaeris genus using the add.species.to.genus function of the phytools package (Revell, 2012) in R version R-4.0.4 (R-Core-Team, 2021). For each pair of added alien and host native species, we calculated the phylogenetic distance using the cophenetic function of the ape package (Paradis & Schliep, 2019) in R. To calculate an overall floral trait distance between each pair of alien and native species, we calculated the Gower dissimilarity (Gower, 1971) based on continuous (flower size) and categorical (flower symmetry, binary floral reflectance components and nectar production) floral traits, using the gowdis function of the FD package (Laliberté et al., 2014; Laliberté & Legendre, 2010) in R. We also calculated single floral trait distances and dissimilarities between the alien and the native species: flower-size distance, flower-symmetry dissimilarity, dissimilarity in each of the four floral reflectance components, and nectar-production-dissimilarity. We calculated absolute and hierarchical floral trait distances (Ferenc & Sheppard, 2020; Kunstler et al., 2012). To assess the association between phylogenetic and floral trait distances, we estimated the strength of the phylogenetic signal, using Pagel’s lambda with the phylosig function of the phytools package for the continuous trait, and the phylogenetic D statistic with the phylo.d function of the caper package (Orme et al., 2012) for the categorical traits.

### Statistical analyses

To test whether flower visitation to alien plant species depended on the phylogenetic and floral trait distances between the alien and the native species, we analyzed the total number of flower visits to the added alien plants, the proportion of flower visits to the alien plant relative to the sum of flower visits to both alien and host native plants, and the similarity between the flower visitor compositions of the alien and the native plants.

We analyzed the total number of flower visits to the alien plants with negative binomial generalized linear mixed models using the glmer.nb function in the lme4 package (Bates et al., 2014) in R. To account for potential variation due to floral characteristics of the observed plants, we included the number of observed flower units of the native plant, the number of observed flower units of the alien plant, the flower size, the flower symmetry (radial=0, bilateral=1), the floral reflectance binary categories Wavelength 401-470 nm, Wavelength 471-540 nm, Wavelength 541-610 nm and Wavelength 611-680 nm of the alien plants, and the nectar production (absent=0, present=1). To account for potential variation of flower-visitor activity during the day, we additionally included the time interval during the day (one-hour intervals as a discrete variable) as an explanatory variable. To test for the effects of either the phylogenetic or the floral trait distance between the alien and the native species, we also included them as explanatory variables. All covariates were centered to means of zero and scaled to standard deviations of one. To test for potential nonlinear effects, we included the quadratic terms for the time interval during the day and phylogenetic or floral trait distance. As the latter were centered and scaled, the non-linear effects test for hump- and U-shaped relationships. We also ran the models with linear terms only, and we present the results of the model with the lowest AIC. To account for non-independence of observations within species, we included identities of the added alien species and the host native species as random factors. Models including site as an additional random factor to account for potential variation due to site characteristics, such as floral abundance, did not converge, as native species and site were largely confounded (Table S1). Similarly, models including date of observation as a random factor to account for potential variation due to change in flower visitor communities along the growing season did not converge. To understand whether the resulting patterns were driven by a particular flower visitor group, we also built similar models in which we used as response variables the number of flower visits to the alien plants by Hymenoptera only and by Diptera only, representing the insect orders that contributed most visits (77.24 % and 10.57 %, respectively). We also built models in which we considered each floral trait distance between the alien and the native species separately, instead of an overall floral trait distance. We considered absolute and hierarchical floral trait distances. Additionally, we built models in which we included both phylogenetic and floral trait distances, instead of separately.

To account for the number of flower visits to native plants, we analyzed the logit transformed proportion of flower visits to the alien plant relative to the sum of flower visits to both alien and host native plants (Warton & Hui, 2011) in linear mixed models using the lmer function of the lme4 package in R. We included the same explanatory variables and random factors as in the above models, except for the number of observed flower units of the alien and the native plants, which we replaced with the logit transformed ratio of the number of flower units of the alien plant divided by the sum of the numbers of flower units of the alien and native plants.

To analyze the similarity of the flower-visitor compositions between the alien and the native plants, we calculated a Bray-Curtis similarity index (one minus Bray-Curtis dissimilarity index) based on the abundance of each flower-visitor group (excluding the unknown groups). We analyzed the logit transformed Bray-Curtis similarity index in linear mixed models using the lmer function of the lme4 package in R. We included the same explanatory variables and random factors as in the above models. From these models, we excluded the 1,051 observations for which the number of flower visits to the alien or to the native plant was zero, leaving 2,017 observations with flower visits. For all models, we reported the marginal and conditional *r*^*2*^ (Nakagawa et al., 2017).

## Results

### Effects of phylogenetic and floral trait distances on flower visitation

The alien plants received significantly more flower visits in the middle of the day around 12:00 (Fig. S2) and the number was higher when more of their flower units were observed (Tables 1 and 2). Moreover, they received more visits when they had larger flowers, when their floral reflectance had local maxima in the yellow wavelength interval 541-610 nm, and when they produced nectar (Tables 1 and 2). The alien plants received significantly fewer flower visits when the number of observed flowers on the paired native plant increased (Tables 1 and 2). Among the floral traits, flower size and symmetry had strong phylogenetic signals (Table 3). We found that the phylogenetic distance between the alien and the native plant had significant nonlinear effects on the total number of flower visits to the alien plant and on the proportion of flower visits to the alien relative to the total number of visits to alien and native plants (Fig. 1a and 1c, Table 1). Flower visitation to aliens was lowest when they had intermediate phylogenetic distances to natives (Fig. 1a and 1c). The floral trait distance between the alien and the native plant also had significant nonlinear effects on the total number of flower visits to the alien plant and on the proportion of flower visits to the alien relative to the total number of visits to alien and native plants (Fig. 1b and 1d, Table 2). The alien plants with intermediate floral trait distances to native plants received the most flower visits (Fig. 1b and 1d). We found qualitatively similar results in the analysis of flower visitation by Hymenoptera only (Table S4), but partly different results in the analysis of flower visitation by Diptera only (Table S5). We also found qualitatively similar results in the analyses including both phylogenetic and floral trait distances (Table S6).

**Table 1.** Results of a negative binomial generalized linear mixed model and a linear mixed model testing how the phylogenetic distance between the alien and the native plants influence the total number of flower visits to the alien plant and the proportion of flower visits to the alien relative to the total number of flower visits to the alien and the native plants (n=3068).

| Response variables | Total number of flower visits to the alien plant | Proportion of flower visits to the alien relative to the sum of flower visits to the alien and native plants |
| --- | --- | --- |
| Fixed terms | Estimate (Standard error) | Estimate (Standard error) |
| Intercept | **1.014 (0.302)** | **-2.018 (0.449)** |
| Number of flower units of the native plant | **-0.133 (0.032)** |  |
| Number of flower units of the alien plant | **0.381 (0.045)** |  |
| Number of flower units of the alien divided by the sum of the number of flower units of the alien and the native plants |  | **1.178 (0.073)** |
| Flower size of the alien plant | **0.482 (0.173)** | **1.016 (0.267)** |
| Flower symmetry of the alien plant | 0.145 (0.322) | 0.509 (0.522) |
| Floral reflectance Wavelength 401-470 nm of the alien plant | -0.554 (0.792) | -0.146 (1.087) |
| Floral reflectance Wavelength 471-540 nm of the alien plant | 2.172 (1.646) | 4.225 (2.258) |
| Floral reflectance Wavelength 541-610 nm of the alien plant | **2.116 (1.017)** | **2.875 (1.414)** |
| Floral reflectance Wavelength 611-680 nm of the alien plant | 0.787 (0.648) | 0.822 (0.902) |
| Nectar production of the alien plant | 2.220 (1.227) | 1.150 (1.642) |
| Time during the day | **-0.067 (0.023)** | **-0.101 (0.041)** |
| Time during the day squared | **-0.059 (0.025)** | **-0.121 (0.047)** |
| Phylogenetic distance between the alien and the native plants | **0.545 (0.097)** | **0.593 (0.173)** |
| Phylogenetic distance between the alien and the native plants squared | **0.212 (0.046)** | **0.266 (0.082)** |
| Random terms | SD | SD |
| Alien species | 1.564 | 2.136 |
| Native species | 0.468 | 1.002 |
| Residuals |  | 2.220 |
| AIC | 16600.400 | 13829.740 |
| Marginal $r^2$ | 0.263 | 0.233 |
| Conditional $r^2$ | 0.832 | 0.640 |

Significant model parameters are highlighted in bold ( $p < 0.05$ ), and marginally significant model parameters are italicized ( $p < 0.1$ ).

**Table 2.** Results of a negative binomial generalized linear mixed model and a linear mixed model testing how the floral trait distance based on floral traits between the alien and the native plants influence the total number of flower visits to the alien plant and the proportion of flower visits to the alien relative to the sum of flower visits to the alien and the native plants (n=3068).

| Response variables | Total number of flower visits to the alien plant | Proportion of flower visits to the alien relative to the sum of flower visits to the alien and native plants |
| --- | --- | --- |
| Fixed terms | Estimate (Standard error) | Estimate (Standard error) |
| Intercept | **1.280 (0.295)** | **-1.507 (0.423)** |
| Number of flower units of the native plant | **-0.156 (0.032)** |  |
| Number of flower units of the alien plant | **0.369 (0.046)** |  |
| Number of flower units of the alien divided by the sum of the number of flower units of the alien and the native plants |  | **1.330 (0.073)** |
| Flower size of the alien plant | **0.563 (0.174)** | **0.986 (0.256)** |
| Flower symmetry of the alien plant | 0.233 (0.328) | 0.538 (0.513) |
| Floral reflectance Wavelength 401-470 nm of the alien plant | -0.480 (0.769) | -0.055 (1.015) |
| Floral reflectance Wavelength 471-540 nm of the alien plant | 1.431 (1.597) | 2.587 (2.133) |
| Floral reflectance Wavelength 541-610 nm of the alien plant | *1.838 (0.992)* | **2.507 (1.333)** |
| Floral reflectance Wavelength 611-680 nm of the alien plant | 0.804 (0.632) | 0.872 (0.847) |
| Nectar production of the alien plant | **2.240 (1.191)** | 1.335 (1.538) |
| Time during the day | **-0.068 (0.023)** | **-0.099 (0.041)** |
| Time during the day squared | **-0.059 (0.026)** | **-0.122 (0.047)** |
| Floral trait distance between the alien and the native plants | -0.008 (0.036) | 0.070 (0.065) |
| Floral trait distance between the alien and the native plants squared | **-0.088 (0.025)** | **-0.266 (0.045)** |
| Random terms | SD | SD |
| Alien species | 1.521 | 1.991 |
| Native species | 0.479 | 1.041 |
| Residuals |  | 2.212 |
| AIC | 16620.000 | 13807.730 |
| Marginal $r^2$ | 0.240 | 0.226 |
| Conditional $r^2$ | 0.818 | 0.619 |

666 Significant model parameters are highlighted in bold ( $p < 0.05$ ), and marginally significant model parameters are italicized ( $p < 0.1$ ).

**Table 3.** Strength of the phylogenetic signals for the floral traits of the alien and native species using Pagel’s lambda (continuous trait) and phylogenetic D values (categorical traits).

| Floral traits | Phylogenetic signal |
| --- | --- |
| Flower size | 0.956<sup>a</sup> |
| Flower symmetry | 0.968<sup>b</sup> |
| Floral reflectance Wavelength 401-470 nm | 0.041<sup>b</sup> |
| Floral reflectance Wavelength 471-540 nm | 0.337<sup>b</sup> |
| Floral reflectance Wavelength 541-610 nm | 0.434<sup>b</sup> |
| Floral reflectance Wavelength 611-680 nm | 0.013<sup>b</sup> |
| Nectar production | 0.198<sup>b</sup> |

<sup>a</sup>A value of 1 indicates that the trait follows a pure Brownian motion model of evolution, <sup>b</sup>the probabilities of phylogenetic D values resulting from Brownian phylogenetic structure are shown.

**Fig. 1.**
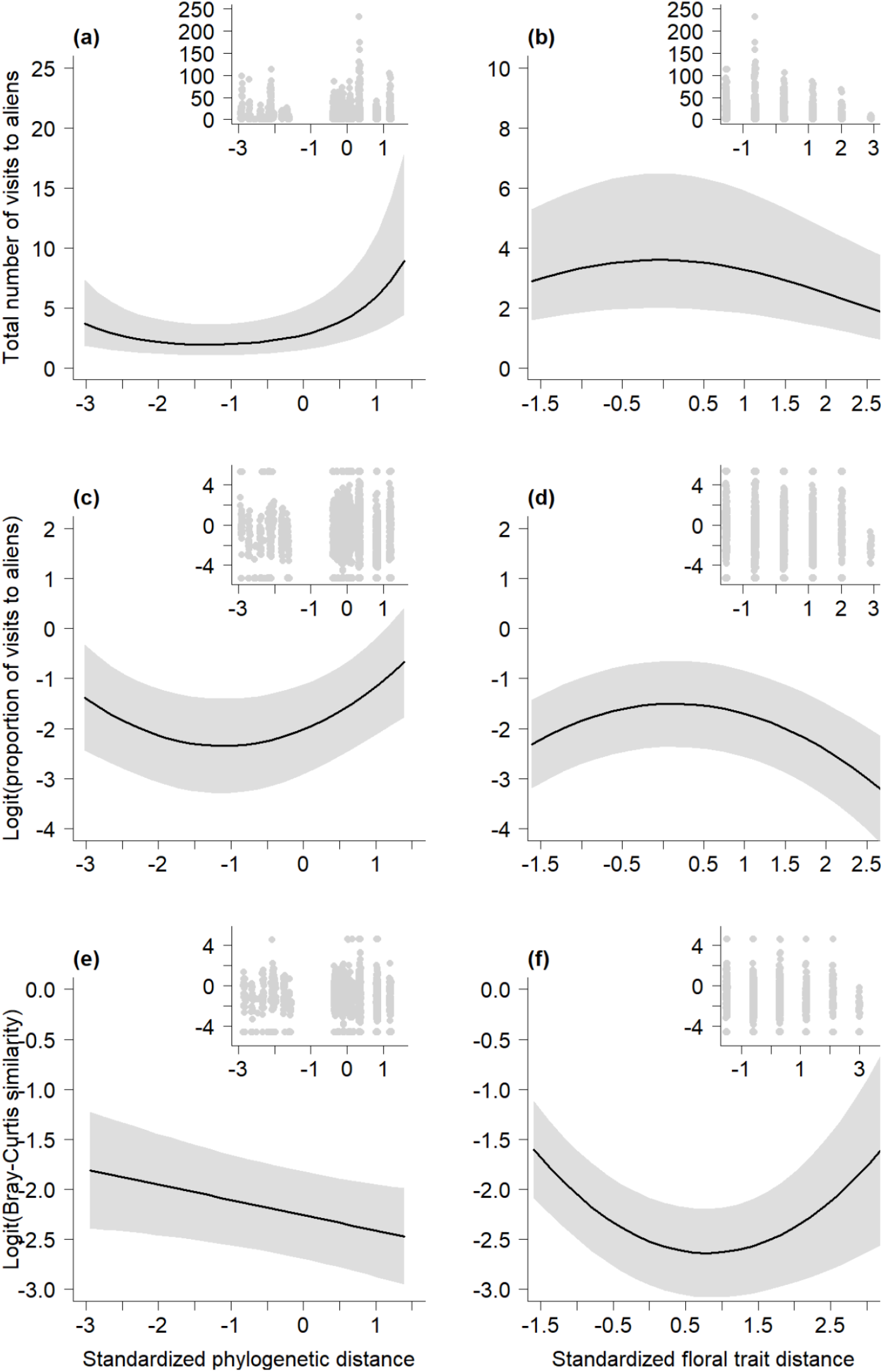
Effects of phylogenetic and floral trait distances on flower visitation to alien plants and on similarity in composition of flower visitors to alien and native plants. (a) Effects of phylogenetic distance on the total number of flower visits to alien plants. (b) Effects of floral trait distance on the total number of flower visits to alien plants. (c) Effects of phylogenetic distance on the proportion of flower visits to alien plants. (d) Effects of floral trait distance on the proportion of flower visits to alien plants. (e) Effects of phylogenetic distance on the similarity in composition of flower visitors to alien and native plants. (f). Effects of floral trait distance on the similarity in composition of flower visitors to alien and native plants. Continuous and dashed curves indicate significant and non-significant effects, respectively. Polygons delimit 95 % confidence intervals. Insets at the right upper corner of each graph show the raw data.

When single absolute floral trait distances were considered separately, we found that alien plants received significantly more flower visits when flower symmetry, the presence of local maxima in the green wavelength interval 471-540 nm of the reflectance spectra and nectar production were dissimilar to the native plants (Table S7). Alien plants received significantly fewer flower visits when flower size distance was larger and when the presence of local maxima in the blue wavelength interval 401-470 nm was dissimilar to the native plants (Table S7). When hierarchical floral trait distances were considered, we found that alien plants received significantly more flower visits when native plants had local maxima in the blue wavelength interval 401-470 nm of the flower reflectance spectra and the alien plants not (Table S8). Alien plants received significantly fewer flower visits when native plants produced nectar and the alien plants not (Table S8).

### Effects of phylogenetic and floral trait distances on similarity in composition of flower visitors to alien and native species

The alien plants received significantly more similar flower visitors to those on native plants when the floral reflectance of the alien plant had local maxima in the yellow wavelength interval 541-610 nm, and marginally significantly when the alien plant produced nectar (Table 4). The alien plants received significantly fewer flower visitors that were similar to those on native plants when the number of observed flowers on the native plants increased (Table 4). We found that the phylogenetic distance between the alien and the native plant had a significant negative effect on the similarity between the flower visitor compositions of the alien and the native plants (Fig. 1e, Table 4). The alien plants with high phylogenetic distances to native plants had the least similar flower visitor composition to native plants. The floral trait distance between the alien and the native plant had a significant nonlinear effect on the similarity between the flower visitor compositions of the alien and the native plants (Fig. 1f, Table 4). The alien plants with intermediate floral trait distances to native plants had the least similar flower visitor composition to native plants (Fig. 1f).

**Table 4.** Results of two linear mixed models testing how the phylogenetic or the floral trait distance based on floral traits between the alien and the native plants influence the similarity between the flower visitor composition of the alien and the native plants (logit Bray-Curtis similarity index, n=2017).

|  | Analysis with phylogenetic<br>distance<br>Estimate (Standard error) | Analysis with floral trait<br>distance<br>Estimate (Standard error) |
| --- | --- | --- |
| Fixed terms |  |  |
| Intercept | **-2.258 (0.219)** | **-2.521 (0.218)** |
| Number of flower units of the native plant | **-0.114 (0.053)** | **-0.102 (0.050)** |
| Number of flower units of the alien plant | 0.002 (0.065) | 0.022 (0.058) |
| Flower size of the alien plant | 0.024 (0.146) | 0.028 (0.172) |
| Flower symmetry of the alien plant | -0.161 (0.344) | 0.016 (0.346) |
| Floral reflectance Wavelength 401-470 nm of the alien plant | 0.242 (0.455) | 0.007 (0.459) |
| Floral reflectance Wavelength 471-540 nm of the alien plant | -0.222 (1.089) | -0.770 (1.110) |
| Floral reflectance Wavelength 541-610 nm of the alien plant | **1.693 (0.650)** | **1.499 (0.655)** |
| Floral reflectance Wavelength 611-680 nm of the alien plant | 0.662 (0.419) | 0.519 (0.419) |
| Nectar production of the alien plant | 1.369 (0.820) | 0.965 (0.826) |
| Time during the day | -0.047 (0.039) | -0.049 (0.040) |
| Time during the day squared | -0.033 (0.043) | -0.030 (0.045) |
| Phylogenetic or floral trait distance between the alien and the native<br>plants | **-0.153 (0.068)** | **-0.290 (0.059)** |
| Phylogenetic or floral trait distance between the alien and the native plants squared |  | **0.180 (0.041)** |
| Random terms | SD | SD |
| Alien species | 0.846 | 0.853 |
| Native species | 0.611 | 0.575 |
| Residuals | 1.704 | 1.689 |
| AIC | 8039.000 | 8008.542 |
| Marginal $r^2$ | 0.045 | 0.062 |
| Conditional $r^2$ | 0.305 | 0.316 |

674 Significant model parameters are highlighted in bold ( $p < 0.05$ ), and marginally significant model parameters are italicized ( $p < 0.1$ ).

## Discussion

In a field experiment simulating invaded co-flowering communities, we found that flower visitation to alien species was highest when they had intermediate floral trait distances to native species, but either low or high phylogenetic distances. This apparent discrepancy may be due to different strengths of phylogenetic signal among traits. The alien plants also received more similar flower visitor groups to natives when they had low phylogenetic and either low or high floral trait distances. The non-linear patterns could be the combined result of facilitation for flower visitation (causing a negative relationship between phylogenetic or floral trait distance and flower visitation to alien species) and competition for flower visitors (causing a positive relationship) (Gallien & Carboni, 2017).

### Non-linear effects of phylogenetic and floral trait distances

Environmental filtering would benefit alien plants that are similar to the native ones, and pollination could be one of the environmental filters. Novel pollinator-mediated alien-native plant interactions can be characterized by a positive influence of co-flowering native plants on the pollination of alien plants (facilitation). Pollinator facilitation operates through different trait-based-effect mechanisms including mimic and magnet effects (Braun & Lortie, 2019). Dominant co-flowering native plants can act as mimic or magnet species that attract pollinators to serve the aliens plants, which have usually left their historical pollinators behind. Pollinator facilitation has been documented in different invaded and non-invaded flowering communities (Bergamo et al., 2020; Ha et al., 2021; Molina-Montenegro et al., 2008; Tur et al., 2016). Some previous studies also provided evidence for pollinator facilitation by alien species to co-flowering natives through the magnet species effect (Groulx & Sargent, 2018; Masters & Emery, 2015; Montero-Castaño & Vilà, 2015; Stiers et al., 2014). The non-linear patterns between flower visitation and similarity of the alien and native species in our study suggest flower visitor facilitation by native species to alien species, at least partially. Nevertheless, more detailed assessments on its effects on plant reproduction should be required to understand the exact processes operating.

The observed patterns of novel pollinator-mediated alien-native plant interactions can also be the outcome of a negative influence of co-flowering native plants on the flower visitation to alien plants (competition). Such competitive interactions are expected to be strongest between plant species that are very similar. The mechanisms of competition for pollinators are complex, including effects of the number of visits on quantity and quality of conspecific pollen received (Mitchell et al., 2009). Alien plants co-occurring and sharing pollinators with one or more dominant flowering natives can compete for pollinator attention, leading to a reduction in flower visitation to the aliens. Pollinator facilitation and competition for pollinators can also act simultaneously in pollinator-mediated alien-native plant interactions (e.g. Bergamo et al., 2018). As we did not quantify visitation to aliens in the absence of natives, we could not quantify facilitation and competition directly. However, as the strength of facilitative and competitive interactions is likely to depend on the dissimilarity of the species, the non-linear patterns we found suggest an interplay of facilitation and competition (Gallien & Carboni, 2017).

### A discrepancy between the effects of phylogenetic and floral trait distances

Most studies on Darwin’s Naturalization Conundrum use phylogenetic distance because it should reflect how functionally dissimilar the species are. This is based on the idea that most traits are phylogenetically conserved. While we found strong phylogenetic signals for flower size and symmetry, the signals were much weaker for spectral reflectance and nectar production, suggesting that phylogenetic distance might not entirely capture functional trait distance. Other studies have also found that flower color is not strongly conserved (Rausher, 2008; Shrestha et al., 2014). On the other hand, in contrast with our results, (Ornelas et al., 2007) found a strong phylogenetic signal in nectar volume and sugar production in their study using 289 species. Remarkably, in our study, while flower visitation was highest at intermediate floral trait distances it was lowest at intermediate phylogenetic distances. This shows that patterns for phylogenetic and trait dissimilarity do not need to be consistent, and may reflect different mechanisms.

Whether co-flowering alien and native plants interact via pollinator facilitation and competition for pollinators depends on the degree of pollinator sharing. In an analysis of 29 plant-pollinator networks, (Vamosi et al., 2014) found that pollinators were more likely to visit closely related species. By analyzing the phylogenetic relatedness among both plants and animals in 36 plant-pollinator and 23 plant-frugivore networks, (Rezende et al., 2007) found that phylogenetically closely related species interacted with a similar set of species. In our study, alien plants with high phylogenetic and intermediate floral trait distances to native plants had the least similar flower-visitor composition to natives (Fig. 1). This suggests that floral trait distances may influence pollinator sharing. Also, the higher visitation of alien plants with intermediate floral trait distances may be largely due to visitation by insects that do not visit the native plants. Nevertheless, future studies should identify flower visitors to more resolved taxonomic levels to more accurately assess the relationships of phylogenetic and floral trait distances with pollinator sharing.

In line with previous findings on pollinator-mediated interactions between alien and native plants (Morales & Traveset, 2009), we found that dissimilarity in floral symmetry was associated with competition for pollinators, as indicated by a positive relationship between floral symmetry dissimilarity and flower visitation to alien plants (Table S7). However, our result on dissimilarity in flower color was partly different from previous studies, as dissimilarity in different components of floral reflectance was associated with either competition or facilitation. For example, we found a negative relationship between dissimilarity in the blue wavelength patterns of petals and flower visitation to alien plants (Table S7). This may be driven by the most abundant flower visitors in our study, the bees (Hymenoptera), which frequently prefer the blue wavelengths (Hsu & Yang, 2012; Razanajatovo et al., 2015). Our findings could thus indicate that similarity in blue wavelength patterns in the petals may increase pollinator facilitation by bees. While previous studies on pollinator-mediated community assembly processes were based on patterns of floral trait distributions within communities (de Jager et al., 2011; Fornoff et al., 2017), by using a manipulative experiment and focusing on pairs of alien and native plants, our results suggest an important role of floral trait similarity in the formation of novel interactions.

In our study, the shapes of the relationships of phylogenetic and floral trait distances with flower visitation to alien plants were contrasting (Fig. 1). The reason for this apparent discrepancy could lay in the floral traits considered in the study and that we may not have measured all relevant traits. If the floral traits are evolutionarily conserved, patterns of trait similarity can be reflected by phylogenetic relatedness (Sargent & Ackerly, 2008). Out of the seven traits included in our study, only flower size and symmetry had strong phylogenetic signals (Table 3). Furthermore, single floral trait distances had different directions of effects, suggesting facilitative, neutral or competitive effects (Table S7). While for the number of visits to aliens, our phylogenetic distance model had the best fit (lowest AIC), interestingly, for the proportional visits and the visitor community similarity, the floral trait distance models had the best fit. Thus, our findings suggest that both phylogenetic and floral trait distances influence pollinator mediated alien-native plant interactions.

### Flower visitation as a proxy for reproductive success

As flower visitors vary in their pollination effectiveness and can even be antagonists, by considering only flower visitation, we cannot be completely certain about which visitors are effective pollinators. Charlebois & Sargent (2017) found a significant relationship between change in flower visitation and change in reproductive success, with a large variability in reproductive success unexplained by change in visitation. They suggested that although flower visitation is not the ideal proxy for reproductive success, it is still very useful (Charlebois & Sargent, 2017). Because seed production and its effect on population growth and invasion should be most important for short-lived self-incompatible alien plants, the ability to attract pollinators in the invaded range might be crucial for such species. Our study included both short- and long-lived species, and self-compatible and self-incompatible species (Table S1), but these life-history characteristics were not related to flower visitation (Tables S9 and S10). Future experiments should assess whether the magnitude of pollen limitation of seed production, and subsequent population dynamics of the alien plants is related to phylogenetic and floral trait distances to co-flowering natives.

## Conclusions

By showing nonlinear effects of phylogenetic and floral trait distances to native species on flower visitation to alien species, this study advances our understanding of how alien plants receive pollination services in the invaded range. Multiple mechanisms and processes including an interplay of pollinator facilitation and competition for pollinators can simultaneously act to engage the formation of novel pollination interactions. We illustrate the importance of considering floral traits in plant community ecology studies to understand major ecological processes such as the formation of novel interactions.

## Acknowledgements

We thank Otmar Ficht and Maximilian Fuchs for taking care of the plants in the botanical garden; Gregor Schmitz for helpful discussions during the planning of the experiment; Lothar Damaschek for facilitating our use of the grasslands around the University of Konstanz; Anna Brumer, Joëlle Clot, Linda Richter and Vanessa Schiwietz for their assistance in the field work; Julia Gutbrod, Hannah Lechner, Elena Werner and V. Schiwietz for their help in analyzing the videos; Ekaterina Mamonova-Stift for her help in measuring floral reflectance. MR was supported by the German Research Foundation DFG (RA 3009/1-1) and an Independent Research Grant from the Zukunftskolleg of the University of Konstanz. FRJ and PRA were supported by the German Academic Exchange Service DAAD.

## Authors’ contributions

MR and MvK designed the study, MR, FRJ and PRA collected the data, MR analyzed the data, MR wrote the first draft of the manuscript and all authors contributed to revisions.

## Data Availability

Should the manuscript be accepted, the data supporting the results will be archived in a public repository, and the data DOI will be included in the article.

