## Supplementary Information for "Flower visitation of alien plants is non-linearly related to phylogenetic and floral similarity to native plants"

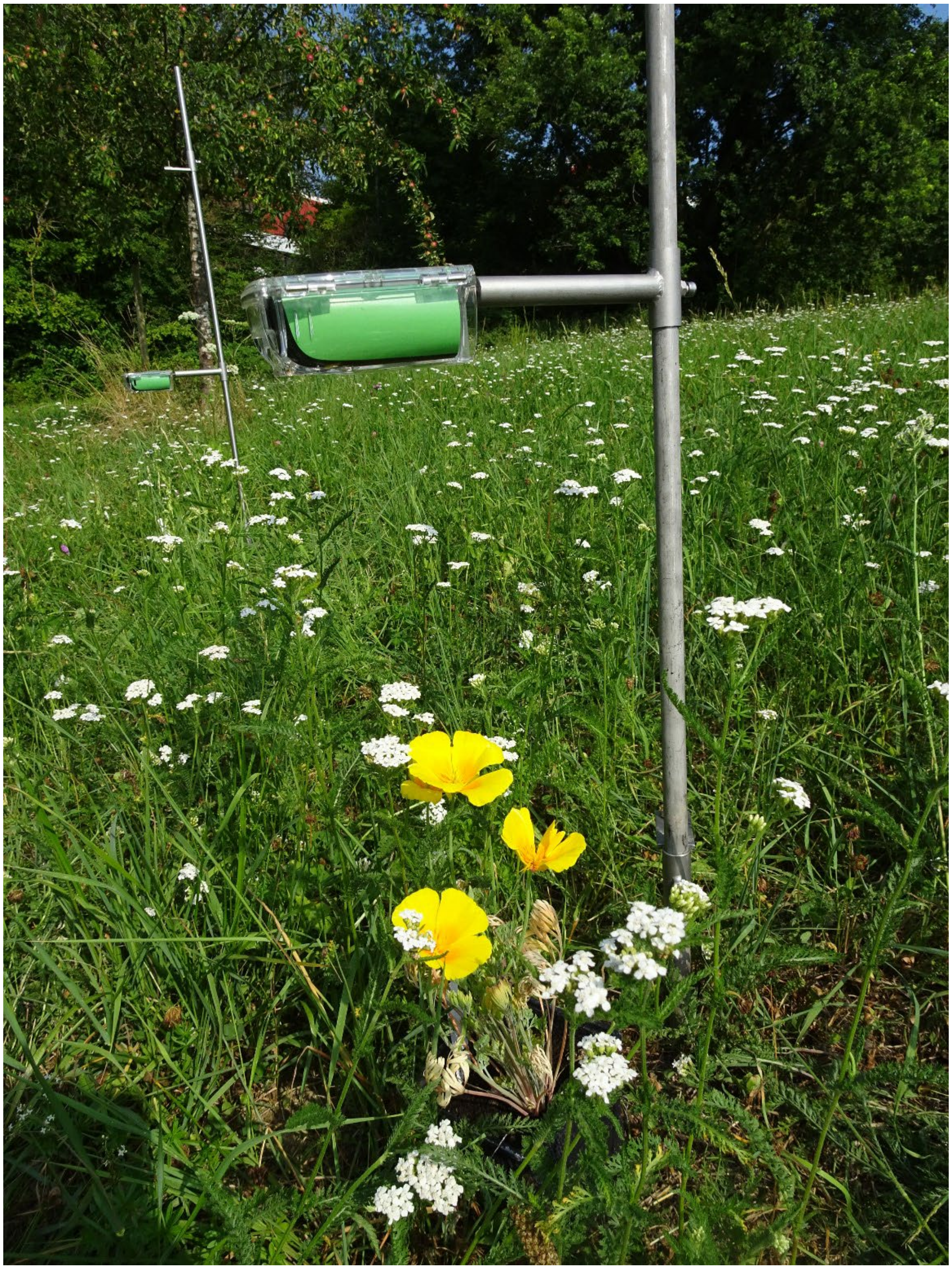

**Fig. S1** Vertical position of the time-lapse-camera at c. 25-30 cm above the observed alien and native flowers.



**Flower visitation of alien plants is non-linearly related to phylogenetic and floral similarity to native plants**

Mialy Razanajatovo, Felana Rakoto Joseph, Princy Rajaonarivelo Andrianina, Mark van Kleunen

**Supplementary information**

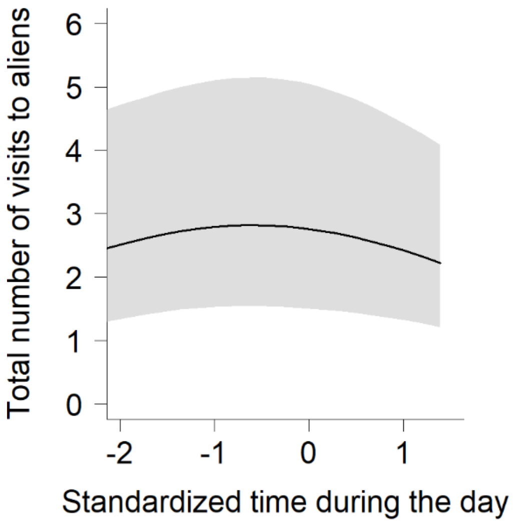


**Fig. S2** A significant nonlinear effect of the time during the day on the total number of visits to added alien flowers. The continuous curve indicates a significant effect. The polygon delimits confidence intervals.

**Supplementary information**

**Table S1** A list of the 34 added alien species used to pair with a host native species to test the effects of phylogenetic and floral trait distances between the alien and the native plants on the flower visitation to the aliens, their life history and breeding system (according to Biolflor except otherwise mentioned), and the sources of plant material.

| **Species** | **Family** | **Life history** | **Breeding system** | **Plant material source** |
|---|---|---|---|---|
| *Ammi majus* | Apiaceae | Annual | Self-compatible | University of Konstanz Botanical garden |
| *Centaurea diffusa* | Asteraceae | Annual | Self-compatible | University of Konstanz Botanical garden |
| *Centaurea nigrescens* | Asteraceae | Perennial | Unknown | Rewisa |
| *Centaurea solstitialis* | Asteraceae | Biennial | Self-incompatible | University of Konstanz Botanical garden |
| *Cerastium tomentosum* | Caryophyllaceae | Perennial | Self-compatible | B&T World Seeds, Jelitto, University of Konstanz Botanical garden |
| *Consolida ajacis* | Ranunculaceae | Annual | Self-compatible | B&T World Seeds |
| *Coriandrum sativum* | Apiaceae | Annual | Self-compatible | Jelitto, University of Konstanz Botanical garden |
| *Crepis setosa* | Asteraceae | Annual | Unknown | University of Konstanz Botanical garden |
| *Datura stramonium* | Solanaceae | Annual | Self-compatible | UFA Samen |
| *Dianthus barbatus* | Caryophyllaceae | Perennial | Self-compatible | Jelitto |
| *Elsholtzia ciliata* | Lamiaceae | Annual | Self-compatible | University of Konstanz Botanical garden |
| *Eschscholzia californica* | Papaveraceae | Annual | Self-incompatible | University of Konstanz Botanical garden, Hof Berggarten |
| *Galinsoga ciliata* | Asteraceae | Annual | Self-compatible | University of Konstanz Botanical garden |
| *Galinsoga parviflora* | Asteraceae | Annual | Self-compatible | University of Konstanz Botanical garden |
| *Hemerocallis fulva* | Xanthorrhoeaceae | Perennial | Self-incompatible | University of Konstanz Botanical garden |
| *Iris versicolor* | Iridaceae | Perennial | Self-incompatible | Ammann |
| *Linaria repens* | Plantaginaceae | Perennial | Self-incompatible | University of Konstanz Botanical garden, wild |
| *Linaria spartea* | Plantaginaceae | Annual | Self-incompatible | Lissabon Ajuda |
| *Lobularia maritima* | Brassicaceae | Annual | Self-incompatible | OBI |

**Supplementary information**

| **Species** | **Family** | **Life history** | **Breeding system** | **Plant material source** |
|---|---|---|---|---|
| *Lychnis coronaria* | Caryophyllaceae | Perennial | Self-compatible | University of Konstanz Botanical garden |
| *Lycopersicon esculentum* | Solanaceae | Annual | Self-compatible | Wyss |
| *Mentha spicata* | Lamiaceae | Perennial | Self-compatible | B&T World Seeds, wild |
| *Nepeta racemosa* | Lamiaceae | Perennial | Unknown | B&T World Seeds |
| *Nicandra physalodes* | Solanaceae | Annual | Self-compatible | University of Konstanz Botanical garden |
| *Nigella damascena* | Ranunculaceae | Annual | Self-compatible | University of Konstanz Botanical garden |
| *Nigella sativa* | Ranunculaceae | Annual[a] | Self-compatible[b] | University of Konstanz Botanical garden |
| *Ornithopus sativus* | Fabaceae | Perennial | Self-compatible | Wyss |
| *Potentilla norvegica* | Rosaceae | Perennial | Pseudogamous apomict[c] | University of Konstanz Botanical garden |
| *Satureja hortensis* | Lamiaceae | Annual | Self-compatible | Jelitto |
| *Sinapis alba* | Brassicaceae | Annual | Self-incompatible | Rieger-Hofmann |
| *Trifolium alexandrinum* | Fabaceae | Annual | Self-compatible | Rieger-Hofmann |
| *Tropaeolum majus* | Tropaeolaceae | Annual | Self-compatible | Rieger-Hofmann |
| *Veronica persica* | Plantaginaceae | Annual | Self-compatible | University of Konstanz Botanical garden |
| *Vicia lutea* | Fabaceae | Annual | Self-compatible | University of Konstanz Botanical garden |

[a]multiple sources (e.g. Ketenoglu et al. 2020), [b]Lloyd 1979, [c]Asker 1970

**Supplementary information**

**Table S2** A list of the 20 host native species used to pair with an added alien species to test the effects of phylogenetic and floral trait distances between the alien and the native plants on the flower visitation to the aliens, the site locations, the density of flowers for the host native species, the density of flowers for all flowering species, and the dates of flower visitation record.

| Species | Family | Sites (latitude, longitude) | Density of flowers (host native species) | Density of flowers (all species) | Dates of flower visitation record |
|---|---|---|---|---|---|
| *Achillea millefolium* | Asteraceae | 47.688132, 9.190459 | 625 | 85 | 20 July 2018 |
| *Centaurea cyanus* | Asteraceae | 47.694388, 9.188946 | 115 | 24 | 29 June 2018 |
| *Centaurea jacea* | Asteraceae | 47.689405, 9.184921<br>47.691026, 9.189862 | 90<br>138 | 30<br>87 | 30 May 2018<br>12 July 2018 |
| *Colchicum autumnale* | Colchicaceae | 47.684417, 9.191214 | 102 | 135 | 16 August 2018 |
| *Daucus carota* | Apiaceae | 47.683169, 9.190011<br>47.68496, 9.190712 | 310<br>589 | 71<br>89 | 13 July 2018<br>27 July 2018 |
| *Filipendula ulmaria* | Rosaceae | 47.6833, 9.190677 | 4140 | 1763 | 20 June 2018 |
| *Hypochaeris radiata* | Asteraceae | 47.682328, 9.19074 | 24 | 10 | 20 September 2018 |
| *Knautia arvensis* | Caprifoliaceae | 47.680987, 9.192175 | 36 | 12 | 6 June 2018 |
| *Leontodon hispidus* | Asteraceae | 47.691111, 9.189951 | 96 | 39 | 9 July 2018 |
| *Leucanthemum vulgare* | Asteraceae | 47.688498, 9.189796 | 200 | 51 | 25 May 2018 |
| *Lotus corniculatus* | Fabaceae | 47.694535, 9.190633<br>47.682927, 9.189491 | 4900<br>1250 | 718<br>212 | 27 June 2018<br>26 July 2018 |
| *Lysimachia vulgaris* | Primulaceae | 47.683908, 9.190606 | 96 | 114 | 15 June 2018 |

**Supplementary information**

| **Species** | **Family** | **Sites (latitude, longitude)** | **Density of flowers (host native species)** | **Density of flowers (all species)** | **Dates of flower visitation record** |
|---|---|---|---|---|---|
| *Lythrum salicaria* | Lythraceae | 47.685954, 9.132723 | 1760 | 627 | 3 August 2018 |
| *Pimpinella saxifraga* | Apiaceae | 47.684922, 9.189618<br>47.694536, 9.190363 | 486<br>468 | 133<br>98 | 28 June 2018<br>4 July 2018 |
| *Prunella vulgaris* | Lamiaceae | 47.694592, 9.193336 | 288 | 79 | 25 July 2018 |
| *Senecio aquaticus* | Asteraceae | 47.687385, 9.192933 | 260 | 119 | 16 July 2018 |
| *Sinapis arvensis* | Brassicaceae | 47.682796, 9.185003 | 84 | 24 | 8 June 2018 |
| *Trifolium pratense* | Fabaceae | 47.694372, 9.192075<br>47.686115, 9.19097 | 584<br>28 | 97<br>22 | 11 July 2018<br>19 September 2018 |
| *Trifolium repens* | Fabaceae | 47.694368, 9.191335 | 714 | 228 | 2 July 2018 |
| *Valeriana officinalis* | Caprifoliaceae | 47.685609, 9.190439 | 4320 | 876 | 14 June 2018 |

**Supplementary information**

**Table S3** Information on nectar production and sources of information for the alien and native species used to test the effects of phylogenetic and floral trait distances on flower visitation of alien species

| Species | Status | Nectar | Source |
|---|---|---|---|
| *Achillea millefolium* | Native | present | Biolflor |
| *Ammi majus* | Alien | present | Biolflor |
| *Anethum graveolens* | Alien | present | Biolflor |
| *Centaurea cyanus* | Native | present | Biolflor |
| *Centaurea diffusa* | Alien | present | Biolflor |
| *Centaurea jacea* | Native | present | Biolflor |
| *Centaurea nigrescens* | Alien | present | Biolflor |
| *Centaurea solstitialis* | Alien | present | Biolflor |
| *Cerastium tomentosum* | Alien | present | Mazzeo et al 2007 |
| *Colchicum autumnale* | Native | present | Biolflor |
| *Consolida ajacis* | Alien | present | Biolflor |
| *Coriandrum sativum* | Alien | present | Biolflor |
| *Crepis setosa* | Alien | present | Mačukanović-Jocić & Jarić 2016 |
| *Datura stramonium* | Alien | present | Biolflor |
| *Daucus carota* | Native | present | Biolflor |
| *Dianthus barbatus* | Alien | present | Biolflor |
| *Elsholtzia ciliata* | Alien | present | Zuo-Dong et al 2006 |
| *Eschscholzia californica* | Alien | absent | Biolflor |
| *Filipendula ulmaria* | Native | absent | Biolflor |
| *Galinsoga ciliata* | Alien | present | Wrzesień & Denisow 2007 |
| *Galinsoga parviflora* | Alien | present | da Silva Camilo et al 2016 |
| *Hemerocallis fulva* | Alien | present | Biolflor |
| *Hypochaeris radiata* | Native | present | Russell 2013 |
| *Iris versicolor* | Alien | present | Biolflor |
| *Knautia arvensis* | Native | present | Biolflor |
| *Leontodon hispidus* | Native | present | Biolflor |
| *Lepidium virginicum* | Alien | present | Tooker et al 2002 |
| *Leucanthemum vulgare* | Native | present | Biolflor |
| *Linaria repens* | Alien | present | Biolflor |
| *Linaria spartea* | Alien | present | Cullen et al 2018 |
| *Lobularia maritima* | Alien | present | Winkler et al 2009 |
| *Lotus corniculatus* | Native | present | Biolflor |
| *Lychnis coronaria* | Alien | present | Witt et al 2013 |
| *Lycopersicon esculentum* | Alien | absent | Bonner & Dickinson 1990 |

| | | | |
|---|---|---|---|
| *Lysimachia vulgaris* | Native | absent | Biolflor |
| *Lythrum salicaria* | Native | present | Biolflor |
| *Mentha spicata* | Alien | present | Biolflor |
| *Nepeta racemosa* | Alien | present | Biolflor |
| *Nicandra physalodes* | Alien | present | Biolflor |
| *Nigella damascena* | Alien | present | Biolflor |
| *Nigella sativa* | Alien | present | Beykaya 2021 |
| *Ornithopus sativus* | Alien | absent | Biolflor |
| *Pimpinella saxifraga* | Native | present | Kapyla 1978 |
| *Potentilla norvegica* | Alien | present | Winkler 2011 |
| *Prunella vulgaris* | Native | present | Biolflor |
| *Salvia aethiopis* | Alien | present | Celep et al 2020 |
| *Satureja hortensis* | Alien | present | Biolflor |
| *Senecio aquaticus* | Native | present | Gottschalk et al 2020 |
| *Sinapis alba* | Alien | present | Biolflor |
| *Sinapis arvensis* | Native | present | Biolflor |
| *Trifolium alexandrinum* | Alien | present | Biolflor |
| *Trifolium pratense* | Native | present | Biolflor |
| *Trifolium repens* | Native | present | Biolflor |
| *Tropaeolum majus* | Alien | present | Biolflor |
| *Valeriana officinalis* | Native | present | Biolflor |
| *Veronica persica* | Alien | present | Biolflor |
| *Vicia lutea* | Alien | present | Biolflor |

**Supplementary information**

**Table S4** Results of two negative binomial generalized linear mixed models testing how the phylogenetic or the floral trait distance between the alien and the native plants influence the number of flower visits by Hymenoptera to the alien plant (n=3068). Significant model parameters are highlighted in bold ($p<0.05$), and marginally significant model parameters are italicized ($p<0.1$).

| | Analysis with phylogenetic distance | Analysis with floral trait distance |
|---|---|---|
| Fixed terms | Estimate (Standard error) | Estimate (Standard error) |
| Intercept | 0.295 (0.390) | 0.604 (0.380) |
| Number of flower units of the native plant | **-0.131 (0.038)** | **-0.157 (0.038)** |
| Number of flower units of the alien plant | **0.416 (0.053)** | **0.392 (0.055)** |
| Flower size of the alien plant | **0.911 (0.204)** | **1.021 (0.206)** |
| Flower symmetry of the alien plant | 0.194 (0.367) | 0.296 (0.376) |
| Floral reflectance Wavelength 401-470 nm of the alien plant | -0.791 (1.006) | -0.695 (0.979) |
| Floral reflectance Wavelength 471-540 nm of the alien plant | 3.206 (2.092) | 2.407 (2.033) |
| Floral reflectance Wavelength 541-610 nm of the alien plant | **3.737 (1.280)** | **3.515 (1.252)** |
| Floral reflectance Wavelength 611-680 nm of the alien plant | 1.032 (0.818) | 1.043 (0.801) |
| Nectar production of the alien plant | 2.397 (1.580) | 2.368 (1.542) |
| Time during the day | -0.010 (0.027) | -0.006 (0.027) |
| Time during the day squared | -0.038 (0.030) | -0.041 (0.030) |
| Phylogenetic or floral trait distance between the alien and the native plants | **0.693 (0.116)** | -0.012 (0.043) |
| Phylogenetic or floral trait distance between the alien and the native plants squared | **0.266 (0.055)** | **-0.079 (0.031)** |
| Random terms | SD | SD |
| Alien species | 1.984 | 1.932 |

| | | |
|---|---|---|
| Native species | 0.643 | 0.636 |
| AIC | 14186.700 | 14216.500 |
| Marginal R-squared | 0.289 | 0.261 |
| Conditional R-squared | 0.876 | 0.863 |

**Table S5** Results of two negative binomial generalized linear mixed models testing how the phylogenetic or the floral trait distance between the alien and the native plants influence the number of flower visits by Diptera to the alien plant (n=3068). Significant model parameters are highlighted in bold (p<0.05), and marginally significant model parameters are italicized (p<0.1).

| | Analysis with phylogenetic distance | Analysis with floral trait distance |
|---|---|---|
| Fixed terms | Estimate (Standard error) | Estimate (Standard error) |
| Intercept | **-2.273 (0.451)** | **-2.105 (0.463)** |
| Number of flower units of the native plant | *-0.122 (0.066)* | **-0.130 (0.065)** |
| Number of flower units of the alien plant | **0.294 (0.068)** | **0.290 (0.067)** |
| Flower size of the alien plant | **-0.680 (0.294)** | **-0.660 (0.290)** |
| Flower symmetry of the alien plant | 0.201 (0.544) | 0.353 (0.558) |
| Floral reflectance Wavelength 401-470 nm of the alien plant | -0.536 (0.686) | -0.551 (0.692) |
| Floral reflectance Wavelength 471-540 nm of the alien plant | -1.748 (1.717) | -2.838 (1.735) |
| Floral reflectance Wavelength 541-610 nm of the alien plant | -0.701 (1.085) | -1.144 (1.071) |
| Floral reflectance Wavelength 611-680 nm of the alien plant | 0.318 (0.638) | -1.068 (1.089) |
| Nectar production of the alien plant | 1.589 (1.189) | 1.584 (1.204) |
| Time during the day | **-0.431 (0.049)** | **-0.434 (0.049)** |
| Time during the day squared | 0.044 (0.056) | 0.042 (0.055) |
| Phylogenetic or floral trait distance between the alien and the native plants | 0.237 (0.220) | **-0.178 (0.078)** |
| Phylogenetic or floral trait distance between the alien and the native plants squared | 0.110 (0.107) | **-0.131 (0.048)** |
| Random terms | SD | SD |
| Alien species | 1.275 | 1.289 |

| | | |
|---|---|---|
| Native species | 1.551 | 1.661 |
| AIC | 5041.100 | 5028.800 |
| Marginal R-squared | 0.100 | 0.103 |
| Conditional R-squared | 0.754 | 0.754 |

**Supplementary information**

**Table S6** Results of a negative binomial generalized linear mixed model and a linear mixed model testing how the phylogenetic and floral trait distances between the alien and the native plants influence the total number of flower visits to the alien plant and the proportion of flower visits to the alien relative to the total number of flower visits to the alien and native plants (n=3068). Significant model parameters are highlighted in bold (p<0.05), and marginally significant model parameters are italicized (p<0.1).

| Response variables | Total number of flower visits to the alien plant | Proportion of flower visits to the alien relative to the sum of flower visits to the alien and native plants |
|---|---|---|
| Fixed terms | Estimate (Standard error) | Estimate (Standard error) |
| Intercept | **1.104 (0.297)** | **-1.746 (0.436)** |
| Number of flower units of the native plant | **-0.136 (0.032)** | |
| Number of flower units of the alien plant | **0.384 (0.045)** | |
| Number of flower units of the alien divided by the sum of the number of flower units of the alien and the native plants | | **1.320 (0.074)** |
| Flower size of the alien plant | **0.401 (0.174)** | **0.757 (0.262)** |
| Flower symmetry of the alien plant | 0.149 (0.324) | 0.352 (0.512) |
| Floral reflectance Wavelength 401-470 nm of the alien plant | -0.575 (0.772) | -0.152 (0.999) |
| Floral reflectance Wavelength 471-540 nm of the alien plant | 1.584 (1.620) | 2.620 (2.102) |
| Floral reflectance Wavelength 541-610 nm of the alien plant | *1.919 (0.995)* | **2.491 (1.313)** |
| Floral reflectance Wavelength 611-680 nm of the alien plant | 0.778 (0.632) | 0.907 (0.835) |
| Nectar production of the alien plant | *2.316 (1.196)* | 1.534 (1.515) |
| Time during the day | **-0.070 (0.023)** | **-0.100 (0.041)** |
| Time during the day squared | **-0.060 (0.026)** | **-0.120 (0.047)** |
| Phylogenetic distance between the alien and the native plants | **0.567 (0.097)** | **0.582 (0.173)** |

**Supplementary information**

| | | |
|---|---|---|
| Phylogenetic distance between the alien and the native plants squared | **0.223 (0.046)** | **0.279 (0.082)** |
| Floral trait distance between the alien and the native plants | -0.027 (0.037) | 0.070 (0.067) |
| Floral trait distance between the alien and the native plants squared | **-0.093 (0.025)** | **-0.267 (0.045)** |
| Random terms | SD | SD |
| Alien species | 1.521 | 1.958 |
| Native species | 0.480 | 1.084 |
| Residuals | | 2.208 |
| AIC | 16589.900 | 13806.380 |
| Marginal R-squared | 0.266 | 0.238 |
| Conditional R-squared | 0.826 | 0.624 |

**Supplementary information**

**Table S7** Results of a negative binomial generalized linear mixed model and a linear mixed model testing how floral trait distances between the alien and the native plants (absolute distances) influence the total number of flower visits to the alien plant and the proportion of flower visits to the alien relative to the sum of flower visits to the alien and native plants (n=3068). Significant model parameters are highlighted in bold (p<0.05), and marginally significant model parameters are italicized (p<0.1).

| | Total number of flower visits to the alien plant | Proportion of flower visits to the alien relative to the sum of flower visits to the alien and native plants |
|---|---|---|
| Fixed terms | Estimate (Standard error) | Estimate (Standard error) |
| Intercept | **1.220 (0.285)** | **-1.675 (0.410)** |
| Number of flower units of the native plant | **-0.169 (0.033)** | |
| Number of flower units of the alien plant | **0.413 (0.048)** | |
| Number of flower units of the alien divided by the sum of the number of flower units of the alien and the native plants | | **1.296 (0.073)** |
| Flower size of the alien plant | **0.724 (0.176)** | **1.366 (0.263)** |
| Flower symmetry of the alien plant | 0.524 (0.354) | 1.053 (0.544) |
| Floral reflectance Wavelength 401-470 nm of the alien plant | -0.776 (0.753) | -0.356 (1.062) |
| Floral reflectance Wavelength 471-540 nm of the alien plant | 2.582 (1.625) | **4.877 (2.286)** |
| Floral reflectance Wavelength 541-610 nm of the alien plant | 1.451 (1.014) | 2.007 (1.473) |
| Floral reflectance Wavelength 611-680 nm of the alien plant | 0.578 (0.622) | 0.490 (0.885) |
| Nectar production of the alien plant | **2.346 (1.184)** | 1.615 (1.618) |
| Time during the day | **-0.066 (0.023)** | **-0.096 (0.041)** |
| Time during the day squared | **-0.046 (0.026)** | **-0.128 (0.047)** |
| Flower size distance | **-0.163 (0.066)** | *-0.159 (0.105)* |

| | | |
|---|---|---|
| Flower symmetry dissimilarity | **0.082 (0.064)** | 0.164 (0.112) |
| Floral reflectance Wavelength 401-470 nm dissimilarity | **-0.487 (0.091)** | **-0.600 (0.160)** |
| Floral reflectance Wavelength 471-540 nm dissimilarity | **0.675 (0.333)** | 0.367 (0.581) |
| Floral reflectance Wavelength 541-610 nm dissimilarity | -0.208 (0.292) | -0.497 (0.495) |
| Floral reflectance Wavelength 611-680 nm dissimilarity | 0.083 (0.055) | *0.155 (0.099)* |
| Nectar production dissimilarity | **0.675 (0.326)** | **1.736 (0.437)** |
| Random terms | SD | SD |
| Alien species | 1.482 | 2.080 |
| Native species | 0.441 | 0.752 |
| Residuals | | 2.217 |
| AIC | 16598.300 | 13818.05 |
| Marginal R-squared | 0.233 | 0.233 |
| Conditional R-squared | 0.809 | 0.616 |

**Table S8** Results of a negative binomial generalized linear mixed model and a linear mixed model testing how floral trait distances between the alien and the native plants (hierarchical distances) influence the total number of flower visits to the alien plant and the proportion of flower visits to the alien relative to the sum of flower visits to the alien and native plants (n=3068). Significant model parameters are highlighted in bold (p<0.05), and marginally significant model parameters are italicized (p<0.1).

| | Total number of flower visits to the alien plant | Proportion of flower visits to the alien relative to the sum of flower visits to the alien and native plants |
|---|---|---|
| Fixed terms | Estimate (Standard error) | Estimate (Standard error) |
| Intercept | **1.215 (0.274)** | **-1.801 (0.426)** |
| Number of flower units of the native plant | **-0.147 (0.032)** | |
| Number of flower units of the alien plant | **0.373 (0.046)** | |
| Number of flower units of the alien divided by the sum of the number of flower units of the alien and the native plants | | **1.293 (0.073)** |
| Flower size of the alien plant | **0.599 (0.173)** | **1.228 (0.276)** |
| Flower symmetry of the alien plant | 0.468 (0.390) | 0.936 (0.753) |
| Floral reflectance Wavelength 401-470 nm of the alien plant | 0.383 (0.778) | -0.059 (1.232) |
| Floral reflectance Wavelength 471-540 nm of the alien plant | 0.916 (1.640) | 3.958 (2.535) |
| Floral reflectance Wavelength 541-610 nm of the alien plant | **2.367 (0.999)** | **2.919 (1.533)** |
| Floral reflectance Wavelength 611-680 nm of the alien plant | 0.998 (0.637) | 0.330 (1.011) |
| Nectar production of the alien plant | 1.424 (1.192) | -0.719 (1.790) |
| Time during the day | **-0.067 (0.023)** | **-0.100 (0.041)** |
| Time during the day squared | **-0.062 (0.026)** | **-0.123 (0.047)** |
| Flower size distance | 0.120 (0.129) | 0.095 (0.214) |

**Supplementary information**

| | | |
|---|---|---|
| Flower symmetry dissimilarity | *0.228 (0.126)* | 0.197 (0.327) |
| Floral reflectance Wavelength 401-470 nm dissimilarity | **0.527 (0.137)** | 0.022 (0.355) |
| Floral reflectance Wavelength 471-540 nm dissimilarity | -0.169 (0.119) | 0.019 (0.291) |
| Floral reflectance Wavelength 541-610 nm dissimilarity | *0.201 (0.105)* | 0.059 (0.273) |
| Floral reflectance Wavelength 611-680 nm dissimilarity | 0.087 (0.139) | -0.359 (0.348) |
| Nectar production dissimilarity | **-0.404 (0.139)** | **-0.842 (0.353)** |
| Random terms | SD | SD |
| Alien species | 1.465 | 2.102 |
| Native species | 0.318 | 0.864 |
| Residuals | | 2.224 |
| AIC | 16626.500 | 13838.740 |
| Marginal R-squared | 0.267 | 0.249 |
| Conditional R-squared | 0.806 | 0.633 |

**Table S9** Results of a negative binomial generalized linear mixed model and a linear mixed model testing how life history (short-lived/long-lived) influences the total number of flower visits to the alien plant and the proportion of flower visits to the alien relative to the sum of flower visits to the alien and native plants (n=3068). Significant model parameters are highlighted in bold (p<0.05).

| | Total number of flower visits to the alien plant | Proportion of flower visits to the alien relative to the sum of flower visits to the alien and native plants |
|---|---|---|
| Fixed terms | Estimate (Standard error) | Estimate (Standard error) |
| Intercept | **1.179 (0.280)** | **-1.891 (0.359)** |
| Life history | -0.081 (0.254) | 0.149 (0.307) |
| Random terms | SD | SD |
| Alien species | 1.513 | 1.812 |
| Native species | 0.423 | 0.757 |
| Residuals | | 2.351 |
| AIC | 16733.200 | 14138.130 |
| Marginal R-squared | 0.002 | 0.002 |
| Conditional R-squared | 0.748 | 0.412 |

**Table S10** Results of a negative binomial generalized linear mixed model and a linear mixed model testing how breeding-system (self-compatible/self-incompatible) influences the total number of flower visits to the alien plant and the proportion of flower visits to the alien relative to the sum of flower visits to the alien and native plants (n=2823). Significant model parameters are highlighted in bold (p<0.05).

| | Total number of flower visits to the alien plant | Proportion of flower visits to the alien relative to the sum of flower visits to the alien and native plants |
|---|---|---|
| Fixed terms | Estimate (Standard error) | Estimate (Standard error) |
| Intercept | **1.100 (0.311)** | **-1.947 (0.387)** |
| Breeding system | 0.239 (0.309) | 0.357 (0.362) |
| Random terms | SD | SD |
| Alien species | 1.574 | 1.836 |
| Native species | 0.420 | 0.767 |
| Residuals | | 2.317 |
| AIC | 15276.500 | 12926.560 |
| Marginal R-squared | 0.016 | 0.013 |
| Conditional R-squared | 0.768 | 0.432 |
